# Circular RNAs accumulate in ageing human placental tissue and in stillbirth, leading to DNA damage and cellular senescence

**DOI:** 10.1101/2024.09.02.610398

**Authors:** Anya L. Arthurs, Matilda R. Jackson, Dylan McCullough, Hamish S. Scott, Christopher P. Barnett, Stuart T. Webb, Melanie D. Smith, Tanja Jankovic-Karasoulos, Gustaaf A. Dekker, Claire T. Roberts

## Abstract

**Background:** Unexplained stillbirth may occur due to premature placental ageing, with unexpected deterioration of placental function for gestational age. Circular RNAs (circRNAs) are enzyme-resistant RNA molecules that accumulate in ageing tissues. Furthermore, circRNAs bind gDNA directly, forming circRNA:DNA complexes which can induce DNA breaks and genomic instability.

**Objectives:** This study investigated tissue ageing and circRNA accumulation with gestational age in healthy and stillbirth placentae, and determined whether circRNAs directly interact with placental DNA causing DNA damage and cellular senescence.

**Study design:** Placenta samples (n=60 term uncomplicated; n=4 unexplained stillbirth, 23, 26, 31, 34 weeks’ gestation) were assessed. Abundance of 7 candidate circRNAs (circ_0009000, circ_0024157, circ_0061017, circ_0036877, circ_0054624, circ_0111277 and circ_0000284), and their linear transcripts, was quantified using qPCR. Total RNA, in the presence and absence of RNase R and RNase H1, was determined using the Qubit fluorimeter. Physical interaction of candidate circRNAs with DNA was confirmed by DNA:RNA ImmunoPrecipitation (DRIP)-qPCR. Telomere length was assessed using real-time PCR. Relative abundance of senescence-associated genes was quantified using qPCR. DNA damage was assessed using an alkaline Comet Assay. Patient-derived trophoblast stem cells (TSCs) differentiation into syncytiotrophoblasts or extravillous trophoblasts was confirmed using immunofluorescence microscopy. The effect of circ_0000284 knockdown was assessed following transfection with either a siRNA (designed to knockdown circ_0000284) or a scrambled siRNA control, at 5, 10 and 20 nM final concentrations using Lipofectamine RNAiMax. Abundance of circRNAs in maternal blood sampled between 15-16 weeks’ gestation (n=12 control, n=6 women who went on to have a stillbirth) was determined using qPCR. Appropriate statistical analyses were undertaken (SPSS).

**Results:** Placental DNA damage, senescence and expression of 7 candidate circRNAs, but not their linear transcripts, were increased in 40 and 41+ weeks’ gestation samples, and in stillbirth, compared with earlier gestations (37-39 weeks’). DRIP-qPCR signal size was significantly larger in term placentae than in enzyme-treated controls, confirming that all candidate circRNA loci bind to placental DNA. Abundance of circRNA was significantly decreased with the addition of RNase H1, compared with all healthy gestation samples, indicating that stillbirth placentae may lack RNase H1. Telomere length is shorter in placentae from stillbirths compared with healthy 37 weeks’ placentae. Depletion of circ_0000284 by specific siRNA in primary cells significantly reduced DNA damage and increased expression of senescence-associated genes compared to control. Abundance of candidate circRNAs are increased in maternal blood at 16 weeks’ gestation for women who went on to have a stillbirth compared with women who had live births.

**Conclusions:** Stillbirth placentae show accelerated ageing with shortened telomeres, premature DNA breaks, increased cellular senescence and accumulation of candidate circRNAs, at levels consistent with older gestation tissue. These circRNAs bind to DNA in the placenta, and circ_0000284 knockdown reduces DNA breaks and senescence in primary placental cells. Therefore, circRNAs play a role in placental ageing and associate with stillbirth, likely via decreased RNase H1 abundance, preventing circRNA degradation and facilitating circRNA accumulation, and subsequent circR-loop formation. circRNAs present a viable method of stillbirth risk screening.

**Graphical Abstract:** 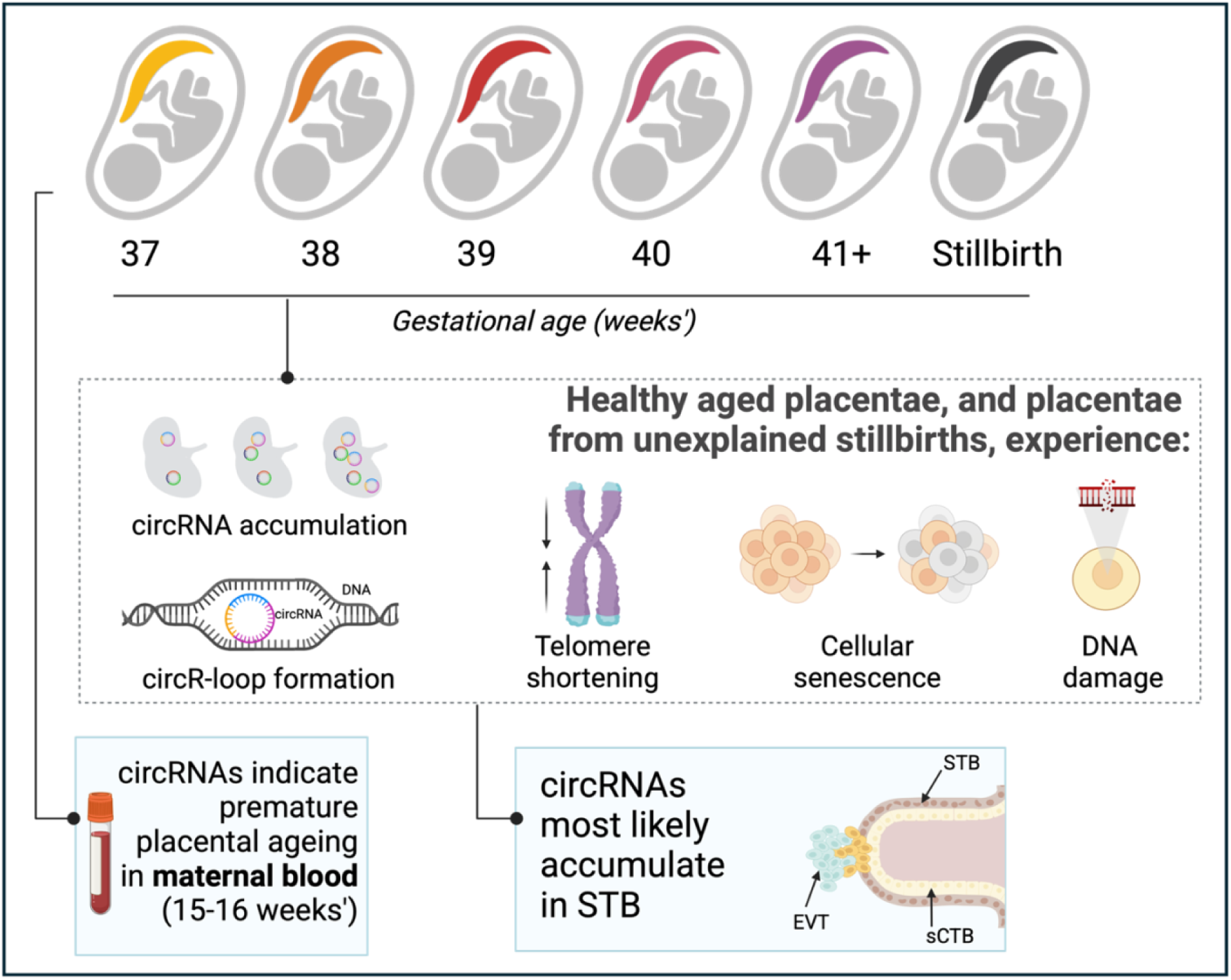

**AJOG at a Glance:**

A. This study was conducted to confirm premature placental ageing in cases of unexplained stillbirth, and to investigate the role of circular RNAs in the process of placental ageing.
B. Circular RNAs accumulate in ageing healthy placenta and accumulate prematurely in stillbirth placentae. Circular RNAs bind directly to placental DNA, inducing DNA breaks and cellular senescence, thereby contributing to overall functional decline of the placenta, reducing its ability to support the fetus. Circular RNAs in maternal blood represents a novel screening tool for detection of stillbirth risk.
C. Premature placental ageing has previously been associated with stillbirth, however circular RNAs have never before been implicated in this process. Circular RNA accumulation in ageing tissues has been shown in model species (i.e. *C. elegans, Drosophila Melanogaster*, mice) but this is the first evidence in human tissues and in the placenta.

## 1. Introduction

Stillbirth is defined as a fetal death prior to birth of a baby of either (i) ≥20 completed weeks’ of gestation or, (ii) ≥400g birthweight^1^. Annually there are 2 million stillbirths worldwide^2^, with ∼2300 occurring every year in Australia, at a rate unchanged for decades^1^. Whilst some cases of stillbirth can be attributed to fetal congenital abnormalities, many cases of stillbirth are “unexplained”, where autopsy that includes fetal genetics does not establish cause. Although some risk factors have been identified^1,3^, the aetiology of unexplained stillbirth is unknown, and is thus impossible to prevent. One theory for the mechanism underpinning unexplained stillbirth is premature placental ageing.

The placenta is a transient organ with a finite life in gestation. Placental ageing is a normal phenomenon resulting in physical changes in late gestation without fetal compromise. Accelerated placental ageing can occur in stillbirth, leading to reduced placental functional capacity for fetal support. Whilst markers of tissue ageing, including measures of telomere length^4^ and DNA damage^3–5^, are reportedly irregularly high in stillbirth placentae, a direct measure of DNA breaks has not previously been reported.

Tissue ageing has been associated with circular RNAs (circRNAs). circRNAs can be generated from most genes and encompass one or multiple exonic or intronic regions of RNA. Importantly, they display tissue-specific patterns of expression, making them unique to their tissue of origin as well as the genome of the individual. circRNAs are inherently stable, hence they can accumulate in tissues with age (previously shown in *Drosophila melanogaster, Caenorhabditis elegans* and murine models^6–8^).

Whilst circRNAs have many functions^9^, of importance is their ability to directly bind to genomic DNA, forming a circRNA:DNA complex (termed a ‘circR-loop’)^10^. Due to the steric hinderance, the presence of a circR-loop can cause RNA polymerase II to stall during transcription, subsequently leading to a double-stranded DNA break. This often results in genomic instability^11^, cellular senescence^12^ and cell death^13^. Whilst their circular form makes them difficult to degrade, RNase H1 can potentially degrade specific circRNAs which form circR-loops, as these maintain a locally open secondary structure^14^ (e.g. ci-ankrd52^15^).

Given that unexplained stillbirth may be due to premature placental ageing, and that circRNAs accumulate in ageing animal tissues, we sought to determine whether circRNAs accumulated with gestational age in healthy and stillbirth placentae. We further investigated whether select circRNAs bind to DNA to form circR-loops and induce DNA breaks. We performed *in vitro* experiments to determine the role of circ_0000284 in contributing to placental cell senescence and DNA damage. Finally, we investigated the potential use of circRNAs as a screening tool for placental ageing culminating in unexplained stillbirth. This study provides novel evidence of circRNA accumulation in ageing human tissue and proposes a potential mechanism for stillbirth pathogenesis.

## 2. Methods

### 2.01 Tissue Samples

*First trimester (6–8 weeks’ gestation) human placentae:* were obtained with informed consent from women undergoing elective termination of pregnancy at the Pregnancy Advisory Centre in the Queen Elizabeth Hospital, in Woodville, South Australia. Ethics approval was obtained from the Central Adelaide Local Health Network Human Research Ethics Committee, HREC/16/TQEH/33, Q20160305. Placentae were collected within minutes of termination upon which villus tissue was washed with Hanks’ Balanced Salt Solution (HBSS; Gibco, Sigma-Aldrich) and transported to the laboratory on ice.

*Healthy placentae from 37-41+ weeks’ gestation:* Term placentae were obtained at the Lyell McEwin Hospital in Elizabeth, South Australia, from women recruited as part of the Screening Tests to predict poor Outcomes of Pregnancy (STOP) (2015–2018) cohort study^16^. The study was registered with the Australian and New Zealand Clinical Trials Registry (ACTRN 12614000985684). All placentae were associated with uncomplicated pregnancies and were collected from white women. Tissue biopsies from placentae were washed in phosphate-buffered saline (PBS) before being snap frozen and held in liquid nitrogen for 15 min, then stored at -80°C. Ethics approval was obtained from the Women’s and Children’s Health Network Human Research Ethics Committee (HREC/14/WCHN/90; STOP). All women provided written informed consent.

*Placentae collected from unexplained stillbirths:* Small placental biopsies from unexplained stillbirth cases were obtained from women who had been recruited in to the NHMRC- and GHFM-MRFF-funded Genomic Autopsy Study^17^. Placental biopsies were obtained as part of the standard autopsy procedure and were stored at -80°C as approved by the Human Ethics Committee of the Women’s and Children’s Health Network, South Australia, Australia (HREC/15/WCHN/35) and Melbourne Health as part of the Australian Genomics Health Alliance protocol (HREC/16/MH/251). Informed consent was obtained to allow access to relevant medical information and subsequent storage of anonymized samples, genomic data and/or medical information in relevant databases and publications.

*Maternal blood samples collected at 15-16 weeks’ gestation:* peripheral non-fasting blood samples were collected as part of the STOP cohort study (see above) and placed on ice prior to processing to isolate plasma. Samples were then stored at -80°C prior to subsequent analyses.

### 2.02 Isolation and cultivation of trophoblast stem cells (TSCs)

TSCs were isolated from n=5 first-trimester placentae as previously described^18,19^. Briefly, first-trimester villus tissue was washed in HBSS (4 °C) prior to manually isolating villus structures for further processing. Villi in HBSS were centrifuged (1000 rpm, 1 min) prior to three consecutive enzymatic digest steps and density gradient centrifugation as per Haider *et al*.^20^. After careful collection of cells between the 35% and 50% Percoll layers, cells were thoroughly washed with HBSS before seeding in fibronectin-coated (20 µg/mL; Millipore) culture plates using a culture medium to promote stemness (media details in **Table 1**). Cells were cultured for 14 days in humidified incubators at 37°C, 1% O_2_ in 5% CO_2_ (to mimic the *in utero* first-trimester placental oxygen tension) prior to differentiation to syncytiotrophoblast or extravillous trophoblast lineages.

**Table 1.**
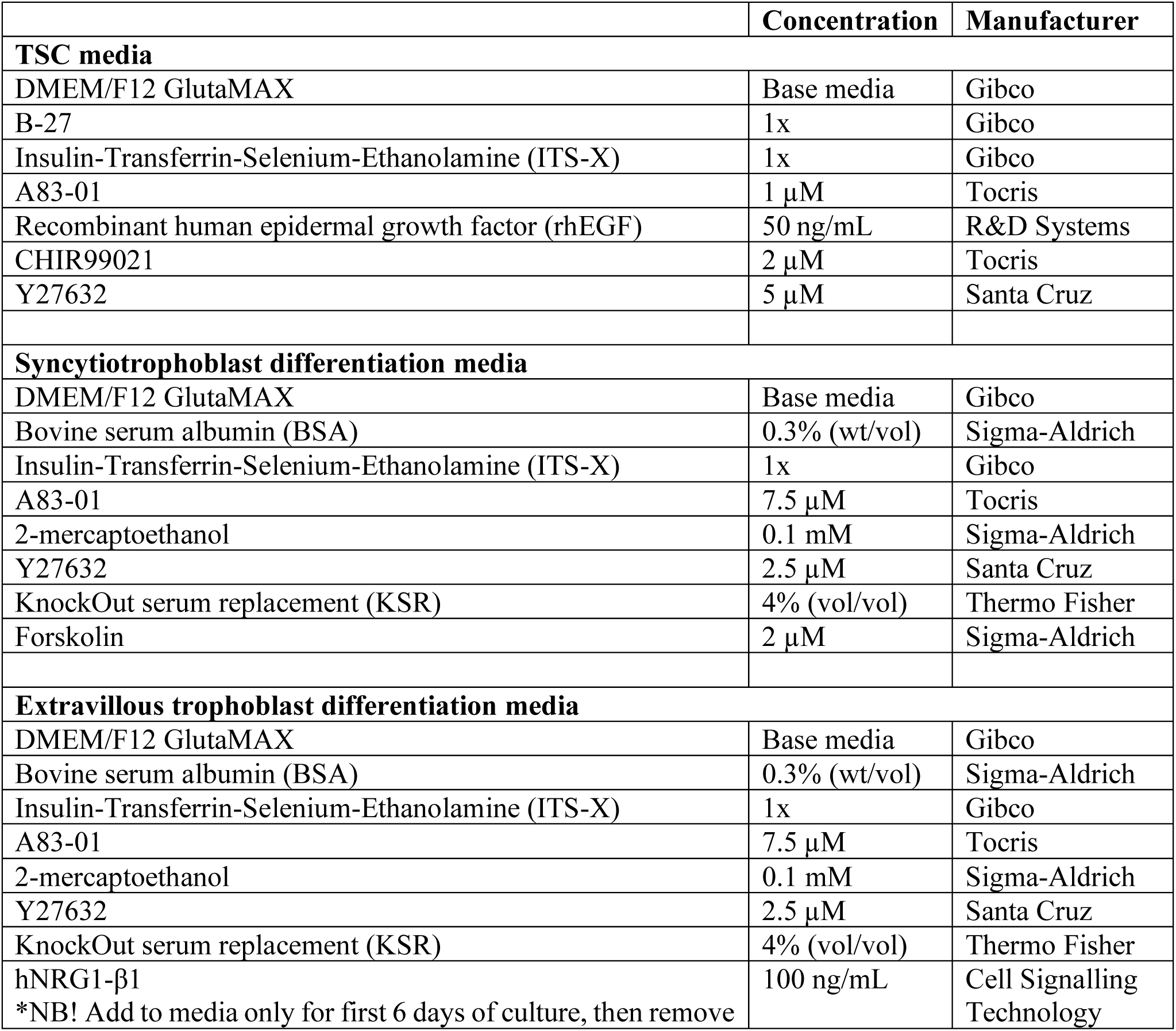
Materials required for preparation of TSC, syncytiotrophoblast and extravillous trophoblast differentiation media.

### 2.03 Differentiation of TSCs into syncytiotrophoblast and extravillous trophoblast lineages

Patient-derived TSCs were reprogrammed to syncytiotrophoblast or extravillous trophoblast lineages using specific culture media as described in Tan *et al.*^21^ (listed in **Table 1**). TSCs were plated in 6-well plates and cultured in syncytiotrophoblast reprogramming media or extravillous trophoblast reprogramming media for a total of four passages. Passaging was completed using TrypLE Express (Gibco) according to manufacturer’s instructions and cells were replated at a density of ∼20% (1:5 dilution) per well. At the time of each passage, a portion of cells were harvested in liquid nitrogen for downstream qPCR analyses. At the end of passage 4 culture, cells were fixed for visualisation or harvested in liquid nitrogen for further experiments.

### 2.04 siRNA transfection of TSCs

TSCs were reprogrammed to syncytiotrophoblasts as above (see **2.03**). siRNA sequence was designed to be complementary to the backsplice junction of circ_0000284 (sequence: ctttattcgagtgattatgattgccatt; scrambled siRNA sequence: tctatcattcatggtatctgtgtatgat). siRNA was produced by Thermo Fisher Scientific (Silencer Select siRNA). Cells were transfected (24-well plates) with the siRNA and scrambled siRNA at 5, 10 and 20 nM final concentrations using Lipofectamine RNAiMAX (Life Technologies) following the manufacturer’s instructions. After optimisation, experiments proceeded with 10 nM final concentration (as the lowest siRNA concentration producing the desired level of knockdown). At 96 hours post-siRNA transfection cells were collected and either (i) immediately used for Comet assay or (ii) snap frozen in liquid nitrogen prior to storage at −80°C for subsequent RNA analyses.

### 2.05 RNA Extraction

*Placenta tissue:* Healthy term and stillbirth placental tissues (25mg) were weighed and washed with phosphate-buffered saline (PBS). Tissue was disrupted by homogenizing for 3.5 min at 30 Hz (TissueLyser, QIAGEN) in 1mL *TRIzol Reagent* (Sigma-Aldrich). Cultured patient-derived cells were suspended in 1mL *TRIzol Reagent*. The manufacturer’s protocol was then followed to isolate total RNA. The purity and integrity of extracted RNA samples were determined using the Qubit 2.0 fluorometer (Thermo Fisher Scientific). Samples from term and stillbirth placenta tissues were then split into two aliquots, with one aliquot to be assessed for circRNA expression levels and the other to be assessed for linear transcript expression levels. Samples to be assessed for circRNA expression levels were heated to 70°C for 5 mins, then immediately cooled below 40°C before incubation with RNase R (10U / reaction; Astral Scientific) for 1 h at 40°C. This step was essential to digest all linear RNAs, leaving only lariat or circular RNA structures.

*Maternal blood:* Total RNA was isolated from 200 μL thawed frozen plasma using the miRNeasy Serum/Plasma kit (Qiagen), according to manufacturer’s instructions and as per Vilades, *et al.*^22^).

### 2.06 RNA Quantification

Total RNA for each sample from term and stillbirth placenta tissues, after incubation of 1 µg of RNA either with RNase R or untreated, was quantified using the Qubit 2.0 fluorimeter (Thermo Fisher Scientific). Total RNA from maternal blood samples was quantified using the Qubit 2.0 fluorimeter as is. As per Drula, *et al.*^23^, Qubit measurement was accomplished with the Qubit™ RNA High Sensitivity (HS) assay kit using 1 µL of the RNA sample, 198 µL of the kit Qubit working solution and 1 µL of the provided fluorophore, as per manufacturer’s instructions. RNA amount was normalised based on the standard curve and input and compared between RNase R treated and untreated groups, in the presence and absence of RNase H1-specific siRNA (as per Wu, *et al.*^24^) per sample to quantify bulk circRNA abundance.

### 2.07 cDNA Synthesis and Quantitative Polymerase Chain Reaction (qPCR)

Synthesis of complementary DNA (cDNA) was conducted beginning with 1μg of total RNA using the QuantiNova Reverse Transcription Kit (QIAGEN) according to the manufacturer’s protocol. qPCR was performed in triplicate and conducted with SYBR Green (QIAGEN) according to manufacturer’s instructions, with *YWHAZ, GAPDH* and *ACTB* as housekeeping genes (all primer sequences in **Table 2**). Custom divergent primers were designed for each circRNA, flanking the backsplice junction, with an amplicon size <300 base pairs. As an additional confirmation of amplification specificity, all primers were assessed using the Primer design BLAST tool^23^. Furthermore, primer specificity was confirmed via melt curve analysis from qRT-PCR. The qPCR conditions were PCR initial activation at 95°C for 5 min followed by 50 cycles of 95°C for 15s, T_m_ (detailed in **Table 2**) for 40s and 72°C for 20s. qPCR results were analysed using the 2-ΔΔCT method^25^. Samples were sorted into groups depending on gestational age of tissue (see **Table 3** for numbers and characteristics of women’s samples used).

**Table 2.**
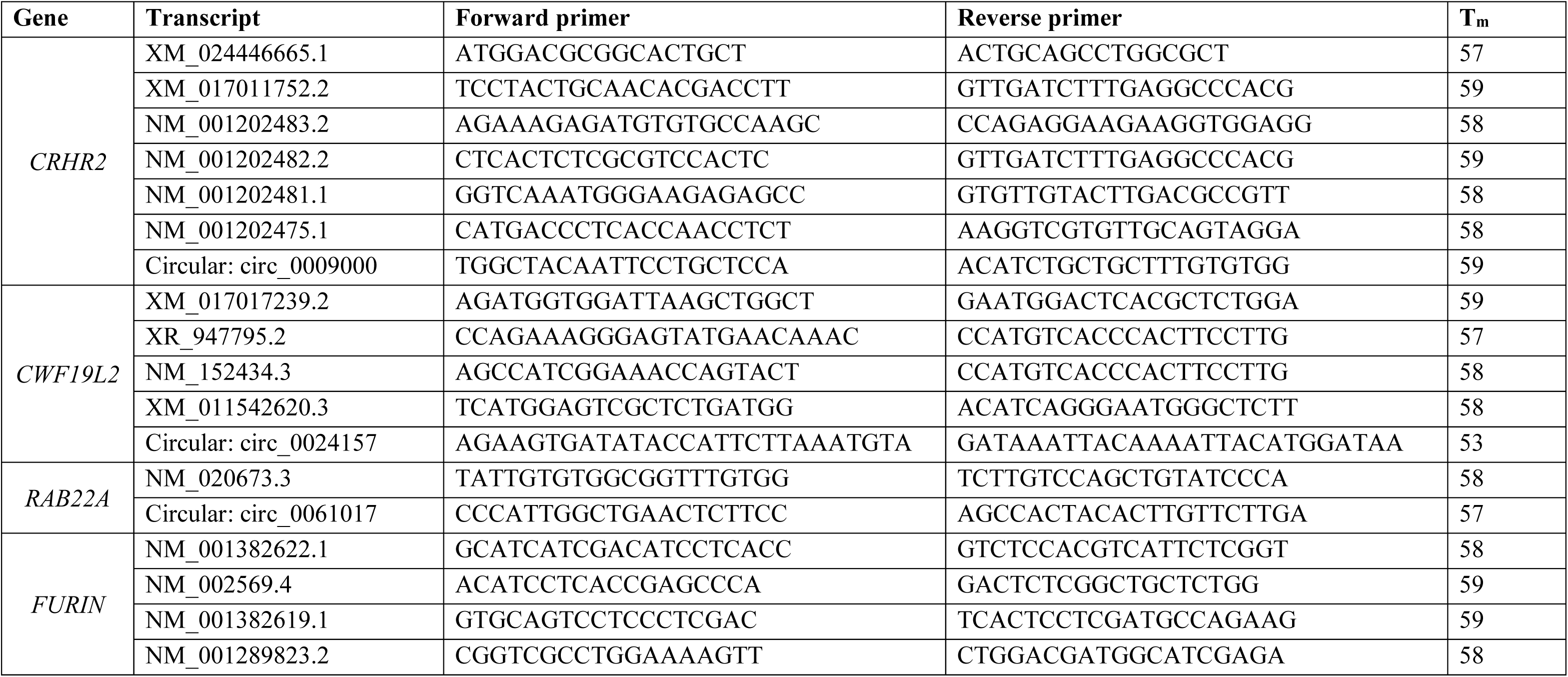
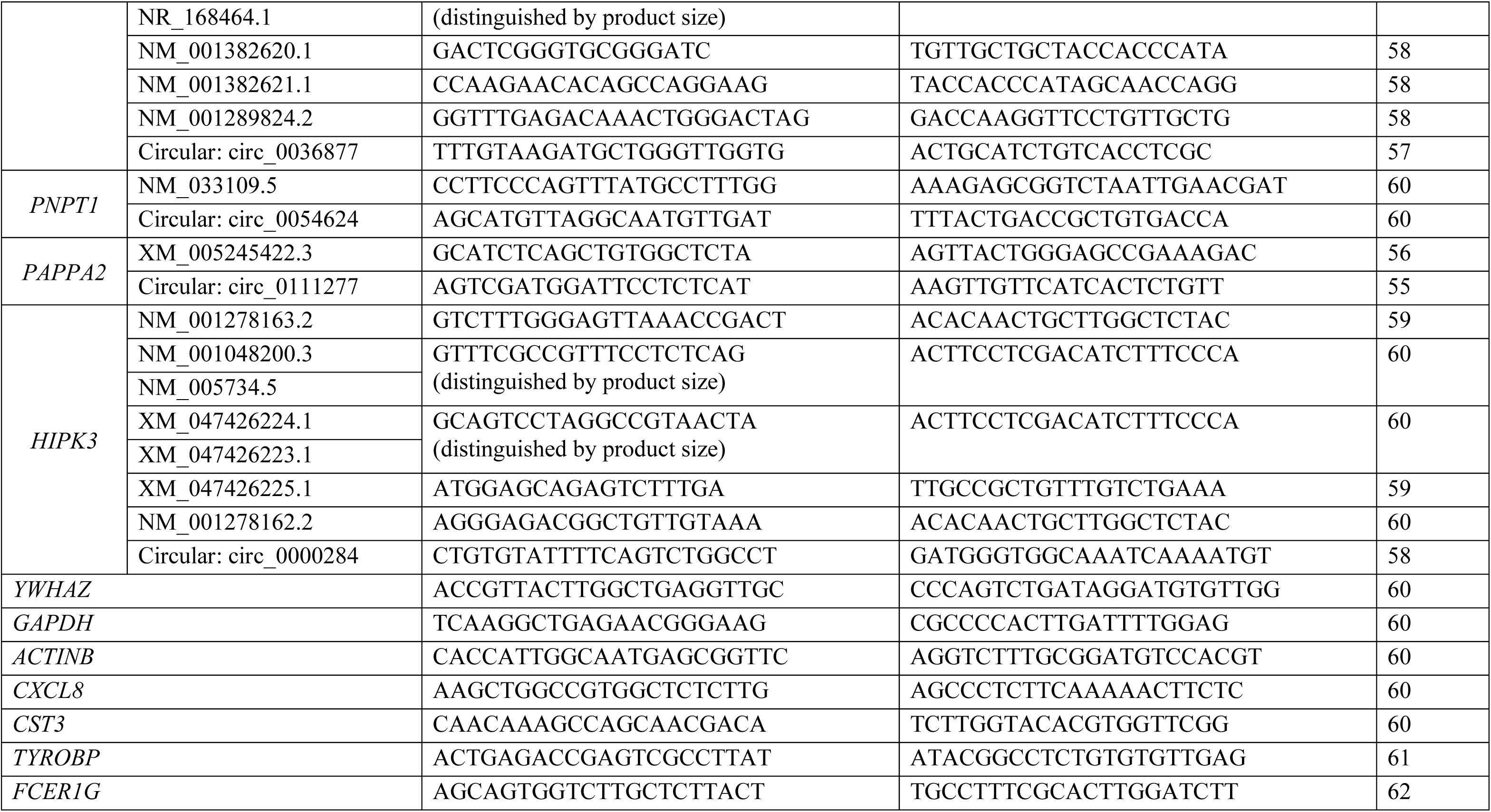
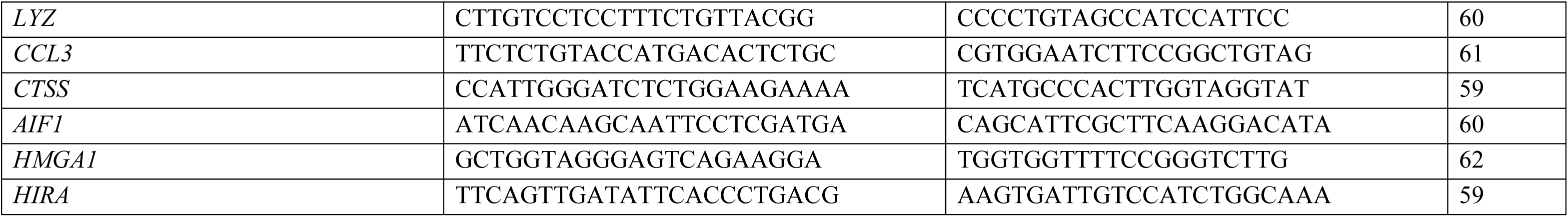
Primer sequences and melt temperatures (T_m_) used for qPCR.

**Table 3.**
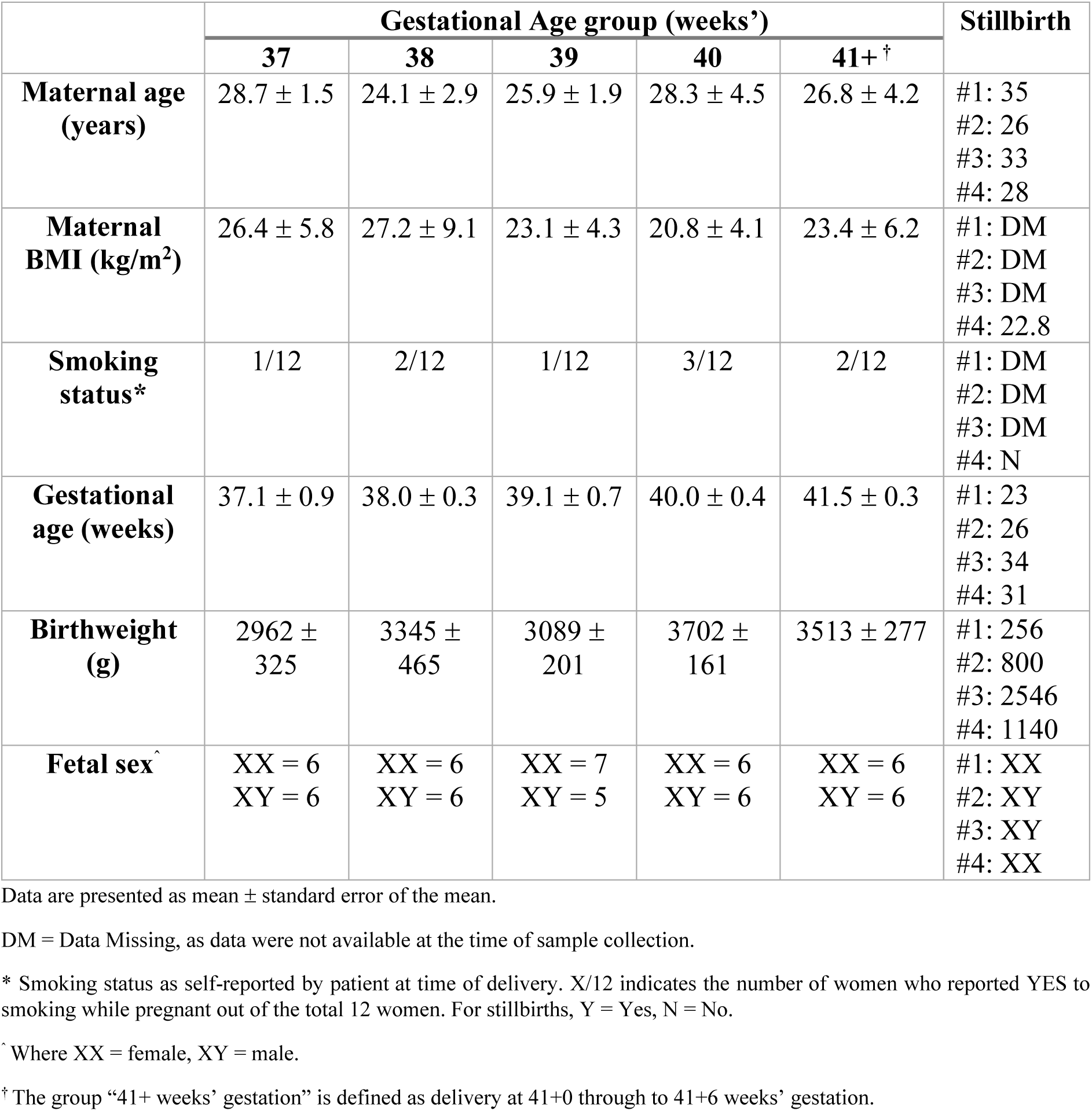
Characteristics of women and infants from the study by gestational age group (weeks’)

|  | Gestational Age group (weeks') |  |  |  |  | Stillbirth |
| --- | --- | --- | --- | --- | --- | --- |
|  | 37 | 38 | 39 | 40 | 41+<sup>†</sup> |  |
| **Maternal age (years)** | 28.7 ± 1.5 | 24.1 ± 2.9 | 25.9 ± 1.9 | 28.3 ± 4.5 | 26.8 ± 4.2 | #1: 35<br>#2: 26<br>#3: 33<br>#4: 28 |
| **Maternal BMI (kg/m<sup>2</sup>)** | 26.4 ± 5.8 | 27.2 ± 9.1 | 23.1 ± 4.3 | 20.8 ± 4.1 | 23.4 ± 6.2 | #1: DM<br>#2: DM<br>#3: DM<br>#4: 22.8 |
| **Smoking status\*** | 1/12 | 2/12 | 1/12 | 3/12 | 2/12 | #1: DM<br>#2: DM<br>#3: DM<br>#4: N |
| **Gestational age (weeks)** | 37.1 ± 0.9 | 38.0 ± 0.3 | 39.1 ± 0.7 | 40.0 ± 0.4 | 41.5 ± 0.3 | #1: 23<br>#2: 26<br>#3: 34<br>#4: 31 |
| **Birthweight (g)** | 2962 ± 325 | 3345 ± 465 | 3089 ± 201 | 3702 ± 161 | 3513 ± 277 | #1: 256<br>#2: 800<br>#3: 2546<br>#4: 1140 |
| **Fetal sex<sup>^</sup>** | XX = 6<br>XY = 6 | XX = 6<br>XY = 6 | XX = 7<br>XY = 5 | XX = 6<br>XY = 6 | XX = 6<br>XY = 6 | #1: XX<br>#2: XY<br>#3: XY<br>#4: XX |

Data are presented as mean ± standard error of the mean.
DM = Data Missing, as data were not available at the time of sample collection.
\* Smoking status as self-reported by patient at time of delivery. X/12 indicates the number of women who reported YES to smoking while pregnant out of the total 12 women. For stillbirths, Y = Yes, N = No.
<sup>^</sup> Where XX = female, XY = male.
<sup>†</sup> The group "41+ weeks' gestation" is defined as delivery at 41+0 through to 41+6 weeks' gestation.

### 2.08 Telomere length analysis

Genomic DNA was extracted from samples using the DNeasy Blood and Tissue Kit (Qiagen) starting with 25 mg tissue. Telomere length was assessed according to the optimised protocol outlined by Cawthon, 2009^26^. Standard curves were generated for telomere lengths from single copy gene amplifications using a reference DNA. The telomere length for each sample was derived based on the ratio of telomere length between the sample and single copy gene standard (*T*/*S* ratio).

### 2.09 Comet Assay

The Abcam Comet Assay Kit (ab238544; Abcam) was used to assess DNA damage in isolated trophoblasts, as per manufacturer’s instructions. For each sample from term and stillbirth placenta tissues, approximately 25 mg tissue (stored at -80°C) was used. For each sample from cultured early gestation patient samples, 2.5x10^5^ cells were used. Alkaline electrophoresis was performed. The DNA integrity of 100 cells per placenta was analysed using CometScore 2.0 software (RexHoover.com). The fluorescence intensity of the comet ‘tail’ was used as a measure of DNA damage (indicated as tail DNA%). Tail DNA was shown as a percentage increase in damage relative to averaged control samples.

### 2.10 DNA:RNA ImmunoPrecipitation (DRIP)

DRIP was performed according to steps 3-25 in the protocol published by Sanz, *et al.* 2019^27^, with some notable exceptions for where tissue was used as a starting material instead of cultured cells. Tissue was first prepared as follows before progressing to step 3 of the protocol.

Approximately 50mg frozen placental villous tissue was submerged in 1mL RLT+ buffer (QIAGEN) with added 0.01% (v/v) β-mercaptoethanol and 0.5% (v/v) Reagent DX (QIAGEN) with 1g beads. Tissue was disrupted by homogenizing for 3.5 min at 30 Hz (TissueLyser, QIAGEN). Cell suspensions were centrifuged (3 min, max speed, 4°C) prior to removal of the supernatant. The cell pellet was washed (sterile DPBS, 5mL) before centrifugation (3 min, 1000 RPM, 4°C) and removal of the supernatant. Steps 3-25 of the protocol^27^ were then followed.

### 2.11 Immunocytochemistry of cultured TSCs

Immunocytochemistry was performed as described by Morosin *et al.*, 2020^28^. Briefly, fixed cells were permeabilised in 0.1% Triton X-100 in PBS (Bio-Rad) before blocking in 1% BSA. Cells were either incubated with E-cadherin primary antibody (Abcam ab1416; syncytiotrophoblast lineage only) or a HLA-G primary antibody (Invitrogen MA1-10359; extravillous trophoblast lineage only) before being washed with PBS and incubated with Alexa Fluor 488 goat anti-mouse IgG (H+L) secondary antibody (Thermofisher, A-11001). Negative controls were included where cells were incubated in the absence of either the primary antibody or both the primary and secondary antibodies. Cells were mounted onto microscope slides using a DAPI mounting medium (ProLong™ Diamond Antifade Mountant with DAPI; Invitrogen). Slides were stored at 4°C and three images were taken per slide (imaged areas were selected at random) using confocal microscopy at 40× magnification.

Syncytialisation was observed where multiple nuclei (blue) existed within one E-cadherin boundary (green). Cells were classed as extravillous trophoblasts where HLA-G was expressed.

### 2.12 Statistical Analysis

Statistical analysis for differences between groups for qPCR and Comet Assay data was undertaken using SPSS Statistics Software. Outliers were removed from the data using a Grubbs’ test. Data were assessed for normality distribution and either a Kruskal-Wallis test or two-way ANOVA tests conducted. Adjustments were made for multiple comparisons (Tukey’s). Differences between groups were considered significant at *p*<0.05.

## 3. Results

### 3.01 Cohort Characteristics

Placenta samples were available from n=60 women who had healthy pregnancies (n=12 per gestational group) and n=4 women who experienced stillbirth. All stillbirths had been assessed by clinical pathologists as “unexplained fetal deaths”, with no noted congenital anomalies.

Maternal and birth characteristics are presented in **Table 3**.

### 3.02 Abundance of candidate circRNAs in placenta across term and in stillbirth

Relative abundance of circ_0009000, expressed as fold change compared to abundance in placentae at 37 weeks’ gestation, was significantly increased in placentae from 40 compared with 37 and 38 weeks’ gestation (*p*_37_=0.0005, *p*_38_=0.0079; **Figure 1A**) and 41+ compared with 37 and 38 weeks’ gestation (all *p*<0.0001), as well as in placentae from stillbirths compared with 37 and 38 weeks’ gestation (*p*_37_<0.0001, *p*_38_=0.0006).

**Figure 1.**
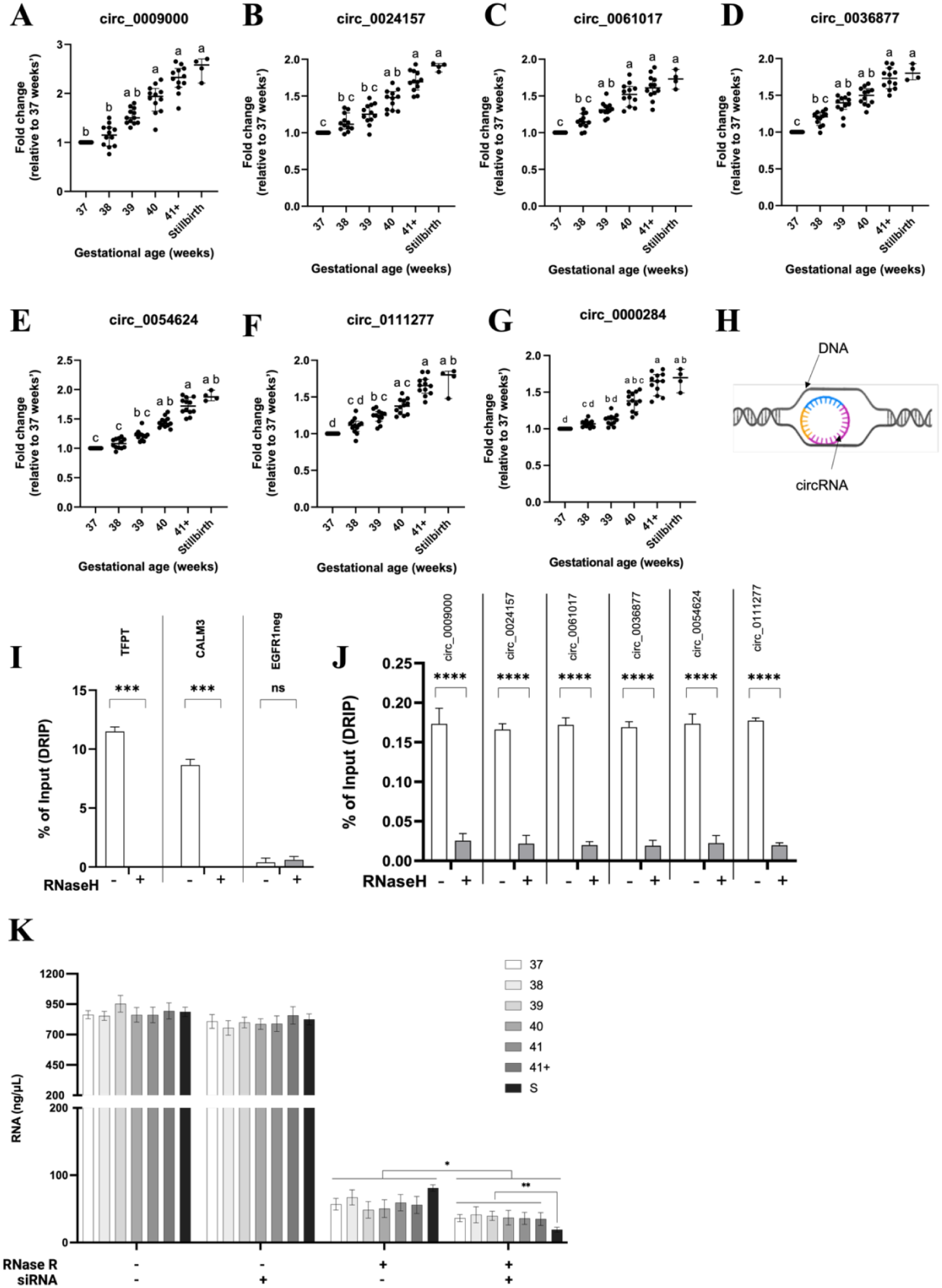
Abundance (as fold change relative to abundance at 37 weeks’ gestation) of **(A)** circ_0009000, **(B)** circ_0024157, **(C)** circ_0061017, **(D)** circ_0036877, **(E)** circ_0054624, **(F)** circ_0111277 and **(G)** circ_0000284, in isolated cells from 37, 38, 39, 40 and 41+ uncomplicated, and stillbirth, placentae. n=12 placentae / uncomplicated gestational group, n=4 stillbirth placentae. Data are presented as scatter plots with bars, and indicate mean ± standard error of the mean. (**H)** Diagram of circR-loop, where the circRNA molecule binds to one strand of helically-unwound DNA. DRIP-qPCR analysis using primers for **(I)** R-loop-positive loci TFPT and CALM3, and R-loop-negative locus EGFR1neg, and **(J)** circ_0009000, circ_0024157, circ_0061017, circ_0036877, circ_0054624, circ_0111277 and circ_0000284. Where indicated, samples were treated with RNase H prior to DRIP. Treatment with RNase H significantly suppressed the DRIP-qPCR signal, consistent with R-loop formation in the tested regions. n=5 placentae with, and n=5 placentae without, RNase H treatment. DRIP-qPCR signal intensity shown as % of Input (DRIP), with indication of mean ± standard error of the mean. **(K)** Quantity of total RNA (ng / μL), starting with ∼1μg total in isolated cells from 37, 38, 39, 40 and 41+ uncomplicated, and stillbirth, placentae. n=12 placentae / uncomplicated gestational group, n=4 stillbirth placentae. RNA quantified in the presence and absence of RNase R (degrades linear transcripts) and an siRNA specific for RNase H1 (enzyme which can degrade circRNAs). Different letters represent statistically significant (p < 0.05) differences. *** p < 0.001, **** p < 0.0001, ns = not significant.

Relative abundance of circ_0024157 was significantly increased in placentae from 40 compared with 37 weeks’ gestation (*p*=0.0003; **Figure 1B**) and 41+ compared with 37, 38 and 39 weeks’ (*p*_37_<0.0001, *p*_38_=0.0002, *p*_39_=0.0124), as well as in placentae from stillbirths compared with 37, 38 and 39 weeks’ gestation (*p*_37_<0.0001, *p*_38_=0.0027, *p*_39_=0.0336).

Relative abundance of circ_0061017 was significantly increased in placentae from 39 compared with 37 weeks’ gestation (*p*=0.0260; **Figure 1C**), 40 compared with 37 and 38 weeks’ gestation (*p*_37_<0.0001, *p*_38_=0.0053) and 41+ compared with 37 and 38 weeks’ gestation (*p*_37_<0.0001, *p*_38_=0.0003), as well as in placentae from stillbirths compared with 37 and 38 weeks’ gestation (*p*_37_<0.0001, *p*_38_=0.0034).

Relative abundance of circ_0036877 was significantly increased in placentae from 39 compared with 37 weeks’ gestation (*p*=0.0151; **Figure 1D**), 40 compared with 37 weeks’ gestation (*p*=0.0002) and 41+ compared with 37 and 38 weeks’ gestation (all *p*<0.0001), as well as in placentae from stillbirths compared with 37 and 38 weeks’ gestation (*p*_37_< 0.0001, *p*_38_=0.0038).

Relative abundance of circ_0054624 was significantly increased in placentae from 40 compared with 37 and 38 weeks’ gestation (*p*_37_<0.0001, *p*_38_=0.0232; **Figure 1E**) and 41+ compared with 37, 38 and 39 weeks’ gestation (*p*_37_<0.0001, *p*_38_<0.0001, *p*_39_=0.0138), as well as in placentae from stillbirths compared with 37 and 38 weeks’ gestation (*p*_37_<0.0001, *p*_38_=0.0012).

Relative abundance of circ_0111277 was significantly increased in placentae from 39 compared with 37 weeks’ gestation (*p*=0.0443; **Figure 1F**), 40 compared with 37 weeks’ gestation (*p*=0.0002) and 41+ compared with 37, 38 and 39 weeks’ gestation (*p*_37_<0.0001, *p*_38_=0.0003, *p*_39_=0.0050), as well as in placentae from stillbirths compared with 37 and 38 weeks’ gestation (*p*_37_<0.0001, *p*_38_=0.0142).

Relative abundance of circ_0000284 was significantly increased in placentae from 40 compared with 37 weeks’ gestation (*p*<0.0001; **Figure 1G**) and 41+ compared with 37, 38 and 39 weeks’ gestation (*p*_37_<0.0001, *p*_38_<0.0001, *p*_39_=0.0158), as well as in placentae from stillbirths compared with 37 and 38 weeks’ gestation (*p*_37_ < 0.0001, *p*_38_ = 0.0042).

### 3.03 Candidate circRNAs bind to DNA to create circR-loops in term placentae

DRIP-qPCR was used to detect circR-loops (DNA:RNA complexes; **Figure 1H**) in pooled placenta samples from gestations 37-41+ weeks’, to determine whether circRNAs which accumulate in the ageing placenta interact directly with DNA.

After completing DRIP (see Methods), DRIP-qPCR analysis was undertaken using either predesigned primers^27^ for known R-loop-positive loci *TFPT* and *CALM3*, and known R-loop-negative locus *EGFR1neg*, or using custom-designed primers (sequences listed in Methods) to target the backsplice junctions of circ_0009000, circ_0024157, circ_0061017, circ_0036877, circ_0054624, circ_0111277 and circ_0000284. All DRIP-qPCR analyses were completed both in samples treated with and without RNase H treatment prior to DRIP. As RNase H cleaves the RNA of RNA:DNA hybrids, it is expected that addition of RNase H to the sample will result in a significantly suppressed R-loop signal and hence, a reduced DRIP-qPCR signal (functioning as a negative control). This was confirmed in our results.

According to Sanz, *et al*.^27^, R-loop positive loci are typically recovered with an efficiency ranging from 1-15% of total input, whereas negative loci are typically recovered with values of <0.1%. This was replicated in our results for placenta tissue (**Figure 1I**), where *TFPT* (mean 11.499%, *p*=0.0008) and *CALM3* (mean 8.644%, *p*=0.0009) produced DRIP-qPCR signals, shown as a % of total DRIP input, significantly larger than their RNase H-treated controls. There was no significant difference between the DRIP-qPCR signals of *EGFR1neg* (mean 0.389%) with and without RNase H treatment.

DRIP-qPCR signal for circ_0009000 (mean 0.173%; **Figure 1J**), circ_0024157 (mean 0.166%), circ_0061017 (mean 0.172%), circ_0036877 (mean 0.169%), circ_0054624 (mean 0.173%), circ_0024157 (mean 0.177%) and circ_0000284 (mean 0.163%) was significantly larger than their RNase H-treated controls (all *p*<0.0001).

### 3.04 circRNAs which accumulate in ageing placenta, and in stillbirth placenta, are subject to RNase H1 degradation

One proposed method for endogenous circRNA degradation is via RNase H1^15^, specifically for circRNAs which can form circR-loops with template DNA due to their locally open secondary structure. RNase H1 cleaves the circR-loop and, in some cases, further degrades the circRNA.

We assessed whether circRNAs in placenta samples from uncomplicated and stillbirth placentae were subject to RNase H1 degradation. Quantity of total RNA for each sample was unchanged with administration of an siRNA which specifically targets RNase H1 (**Figure 1K**). For samples taken from uncomplicated pregnancies delivering at 37, 38, 39, 40 and 41+ weeks’, and in stillbirth placentae, quantity of circRNA (after total RNA treatment with RNase R) was significantly decreased with administration of the RNase H1-specific siRNA (*p*<0.05). Furthermore, quantity of circRNAs was significantly lower after administration of RNase H1-specific siRNA in placentae from stillbirths compared with all other gestations (*p_37_*=0.0094; *p_38_*=0.0087; *p_39_*=0.0072; *p_40_*=0.0090; *p_41+_*=0.0099), indicating that circRNAs from unexplained stillbirth placentae are more susceptible to RNase H1 degradation, likely indicating a deficiency of endogenous RNase H1 in these samples.

### 3.05 Telomere length is decreased in stillbirth compared with healthy term placenta tissue

Relative telomere lengths (**Figure 2A**) were significantly reduced by ∼40% in stillbirth (3.108±1.954) compared with healthy placentae from 37 weeks’ (5.881±2.823; *p* < 0.05). There were no other significant differences in relative telomere lengths between 37, 38 (5.321±2.390), 39 (5.119±2.943), 40 (4.794±2.510), 41+ (4.564±2.428) weeks’ gestation, and stillbirth.

**Figure 2.**
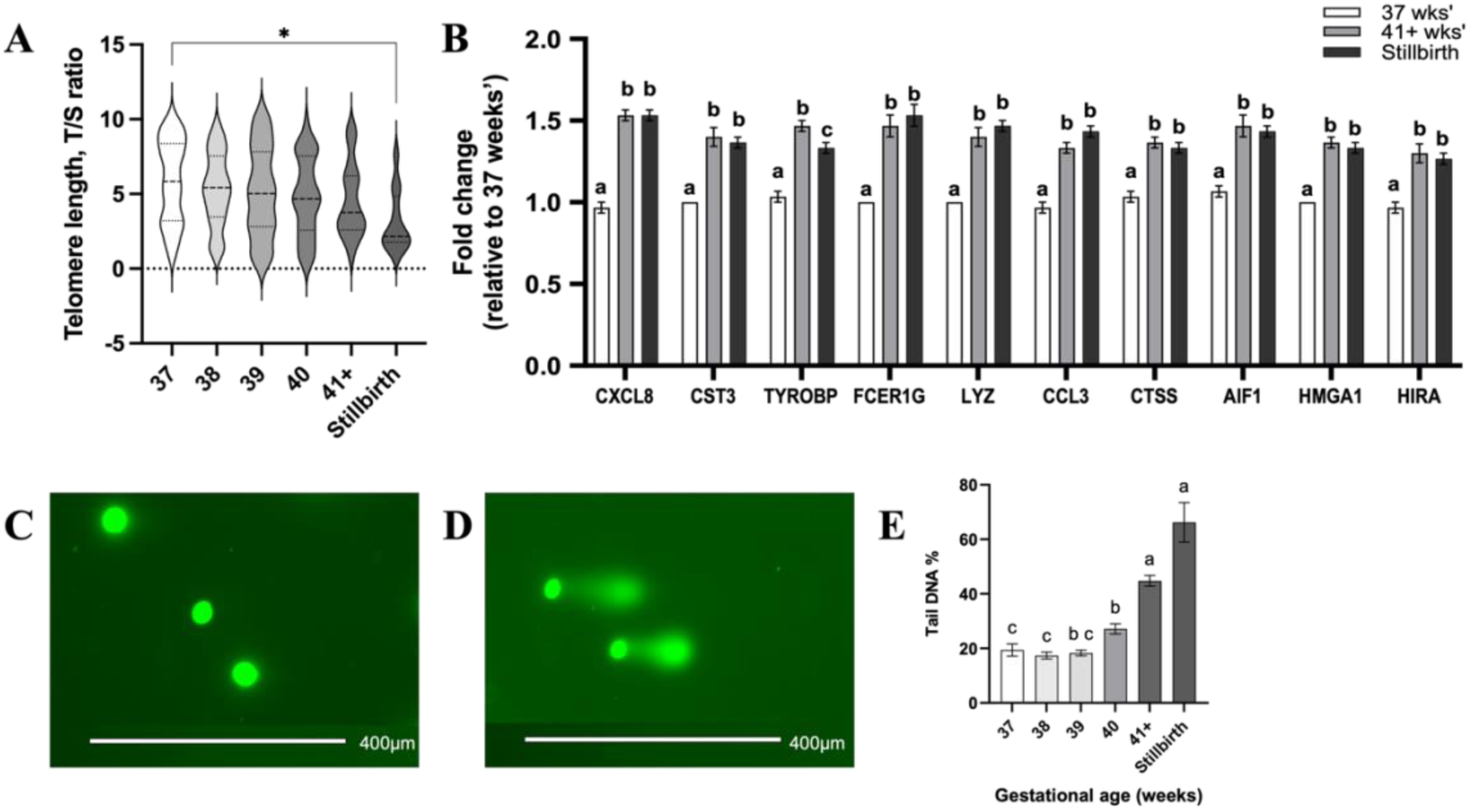
**(A)**Telomere length (expressed as “T/S ratio”, i.e. quantity of telomere DNA divided by quantity of a single copy gene), and **(B)** fold change mRNA expression of senescence-associated genes in placenta tissue from 37 weeks’ gestation uncomplicated pregnancy (white), 41+ weeks’ gestation uncomplicated pregnancy (grey) and stillbirth (black). Data are presented as (A) violin plots and (B) mean ± standard error of the mean. Representative images of **(C)** healthy cells and **(D)** DNA damaged cells, as imaged using epifluorescent microscopy after performing a Comet Assay. **(E)** Levels of DNA damage, as assessed by Tail DNA %, in isolated cells from 37, 38, 39, 40 and 41+ uncomplicated, and stillbirth, placentae. n=12 placentae / uncomplicated gestational group, n=4 stillbirth placentae. 100 cells analysed following Comet Assay, per placenta. Data are presented as mean ± standard error of the mean. Different letters represent statistically significant (p < 0.05) differences. * p < 0.05.

### 3.06 Expression of senescence-associated genes is increased in aged healthy and stillbirth placenta tissue

Saul *et al.*^29^ validated a gene set which identifies senescent cells and predicts senescence-associated pathways in tissue. mRNA expression of senescence-associated genes (**Figure 2B**) was significantly increased in 41+ weeks’ gestation healthy placentae (all *p*<0.0001) and stillbirth placentae (*p*_CXCL8_<0.0001; *p*_CST3_<0.0001; *p*_TYROBP_=0.0003; *p*_FCER1G_<0.0001; *p*_LYZ_<0.0001; *p*_CCL3_<0.0001; *p*_CTSS_<0.0001; *p*_AIF1_<0.0001; *p*_HMGA1_<0.0001; *p*_HIRA_<0.0001) compared with 37 weeks’ gestation healthy placentae.

mRNA expression of *TYROBP* only was significantly increased in 41+ weeks’ gestation healthy placentae compared with stillbirth placentae (*p* = 0.0038).

### 3.07 DNA damage is increased in aged healthy and stillbirth placenta tissue

DNA damage was assessed using the Comet Assay that yields a Tail DNA% metric, an indication of the level of DNA damage in placental cells. Representative images of healthy cells (**Figure 2C**) and DNA damaged cells (**Figure 2D**) are included for reference. Tail DNA % was significantly higher in cells from placentae sampled at 40 compared with 37 and 38 weeks’ gestation (*p*_37_=0.0381, *p*_38_=0.0203; **Figure 2E**), and 41+ compared with 37, 38, 39 and 40 weeks’ gestation (all *p*<0.0001) as well as in placentae from stillbirths compared with 37, 38, 39 and 40 weeks’ gestation (*p*_37_<0.0001, *p*_38_<0.0001, *p*_39_<0.0001, *p*_40_=0.0001).

### 3.08 circ_0000284 is primarily expressed in syncytiotrophoblasts

Patient-derived TSCs (**Figure 3A**) were isolated and underwent cellular reprogramming to either syncytiotrophoblasts (**Figure 3B**) or extravillous trophoblasts (**Figure 3D**). qPCR detected circ_0000284 in syncytiotrophoblasts (**Figure 3C**), where abundance increased with each passage (mean P1=0.045, P2=0.064, P3=0.098, P4=0.120), but was not detected in extravillous trophoblasts (**Figure 3E**).

**Figure 3.**
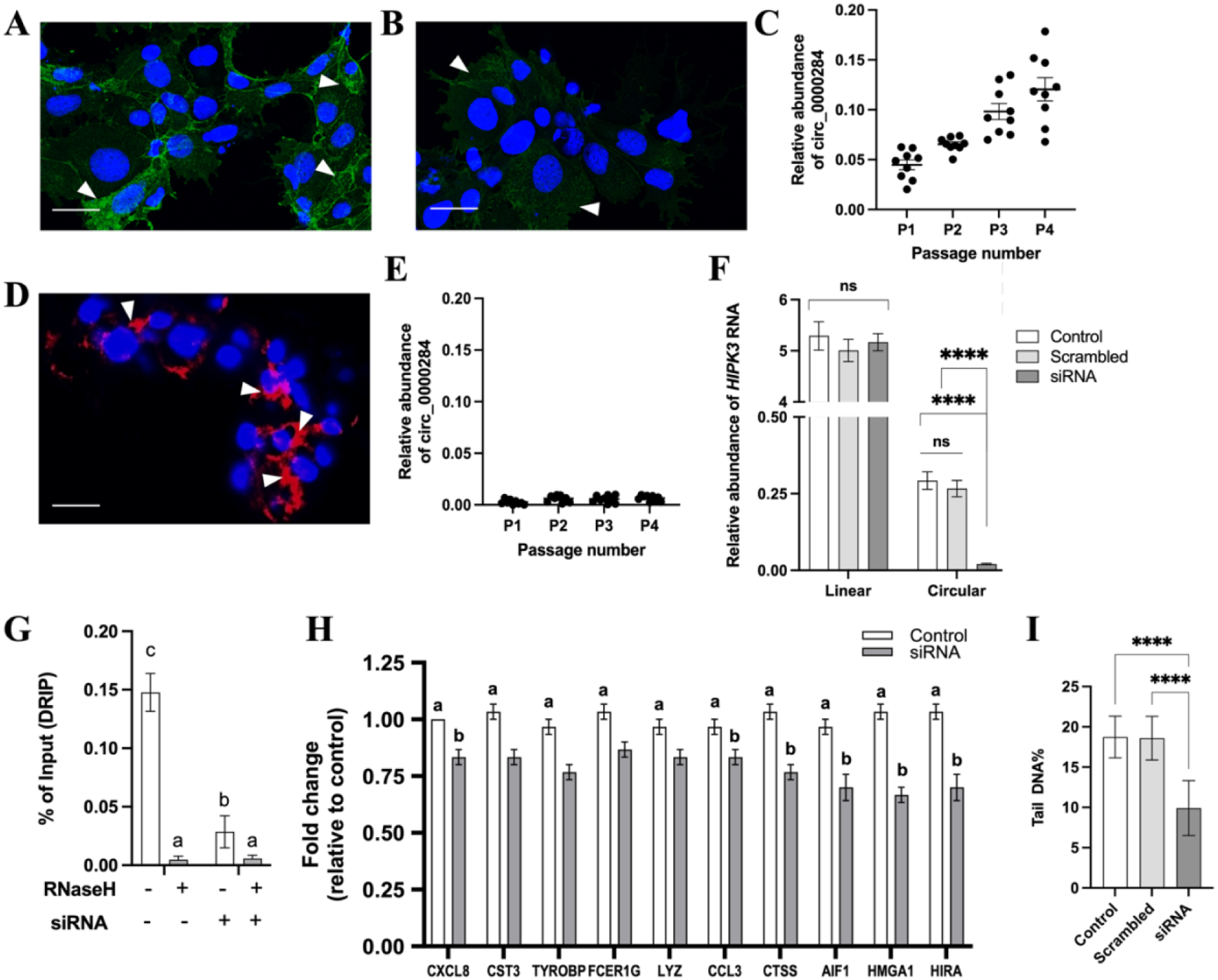
Representative immunocytochemistry images of **(A)** not syncytialised TSCs, **(B)** syncytialised TSCs (i.e. reprogrammed to syncytiotrophoblasts) and **(D)** TSCs reprogrammed to extravillous trophoblasts. Scale bars: 10 µm. Nuclei are stained with DAPI (blue), green stain is for E-cadherin (cell membrane), red stain is for HLA-G (expressed in extravillous trophoblasts). White arrows indicate E-cadherin staining (A, B) and HLA-G staining (D). Relative abundance of circ_0000284 in cultured **(C)** syncytiotrophoblasts, and **(E)** extravillous trophoblasts, over four culture passages. Data are presented as scatter plots with bars, and indicate mean ± standard error of the mean. **(F)** Abundance of HIPK3 RNA (“linear”) and circ_0000284 (“circular”) in cultured syncytiotrophoblast cells, either untransfected (“control”), transfected with a scrambled siRNA (“scrambled”) or transfected with an siRNA specific to circ_0000284 (“siRNA”). **(G)** DRIP-qPCR using primers for circ_0000284, in syncytiotrophoblasts treated with combinations of RNase H and siRNA as indicated. **(H)** Fold change mRNA expression of senescence-associated genes in syncytiotrophoblasts, either untreated (white) or treated with siRNA (grey). **(I)** Levels of DNA damage, as assessed by Tail DNA %, in syncytiotrophoblast cells. n=5 patients / group. 100 cells analysed following Comet Assay, per sample. Data are presented as mean ± standard error of the mean. Different letters represent statistically significant (p < 0.05) differences. **** p < 0.0001, ns = not significant.

### 3.09 *In vitro* knockdown of circ_0000284

Hallmarks of cellular ageing include cellular senescence and DNA damage^30^. circR-loops introduce steric hinderance with the formation of the circRNA:DNA hybrid, which can promote transcriptional pausing and subsequent DNA breaks^31–33^, resulting in genomic instability^34^. As such, we investigated whether circ_0000284 induces cellular senescence and DNA damage. This was confirmed using a custom-designed circ_0000284-specific siRNA in patient-derived TSCs, reprogrammed to syncytiotrophoblasts.

#### Validation of the siRNA knockdown

Cultured primary syncytiotrophoblasts were either untreated (control), treated with a scrambled siRNA or with a custom-designed siRNA specific to circ_0000284. Cells were then cultured for 72 hours prior to harvesting for qPCR analyses as well as Comet Assay.

As shown in **Figure 3F**, transfection with the scrambled siRNA did not alter the expression of the linear *HIPK3* mRNA (shown as abundance relative to *YWHAZ* and *ACTB*) compared to control. Importantly, administration of the siRNA specific to circ_0000284 did not alter the expression of linear *HIPK3* mRNA. However, the siRNA successfully reduced the relative abundance of circ_0000284 compared with both the untreated cells (control) and cells treated with the scrambled siRNA (all *p*<0.0001).

DRIP-qPCR confirmed that circ_0000284 forms a circR-loop in primary syncytiotrophoblasts (mean 0.148%; *p*<0.0001 compared with RNase H-treated control; **Figure 3G**) and administration of the siRNA specific to circ_0000284 significantly reduced circR-loop formation (mean 0.029%; *p*<0.0001).

### 3.10 Expression of senescence-associated genes is attenuated with depletion of circ_0000284

mRNA expression of senescence-associated genes (**Figure 3H**) was significantly decreased in cells transfected with the siRNA specific to circ_0000284 compared to control; *CXCL8* (*p*=0.0053), *CCL3* (*p*=0.0291), *CTSS* (*p*<0.0001), *AIF1* (*p*<0.0001), *HMGA1* (*p*<0.0001) and *HIRA* (*p*=0.0005). mRNA expression of *CST3*, *TYROBP*, *FCER1G* and *LYZ* was not significantly changed.

### 3.11 DNA damage is mitigated with depletion of circ_0000284

Transfection with the siRNA specific to circ_0000284 in primary syncytiotrophoblasts significantly reduced the level of DNA damage (tail DNA%; **Figure 3I**) compared with both the control (reduced by 41.3%) and scrambled (reduced by 39.5%) groups (all *p*<0.0001). Transfection with the scrambled siRNA did not alter the level of DNA damage (tail DNA%) compared with the control, indicating that depletion of circ_0000284 results in reduced DNA damage.

### 3.12 circRNAs can be detected in blood as a proxy measurement of placental ageing

Candidate circRNAs hsa_circ_0009000 (*p*=0.0137), hsa_circ_0036877 (*p*=0.0002), hsa_circ_011277 (*p*=0.0052) and hsa_circ_0054624 (*p*=0.0158) were significantly more abundant (**Figure 4**) in maternal blood sampled at 15-16 weeks’ gestation in women who went on to have a stillbirth compared with women who delivered live babies. Abundance of hsa_circ_0024157, hsa_circ_0061017 and hsa_circ_0000284 were not significantly changed between groups.

**Figure 4.**
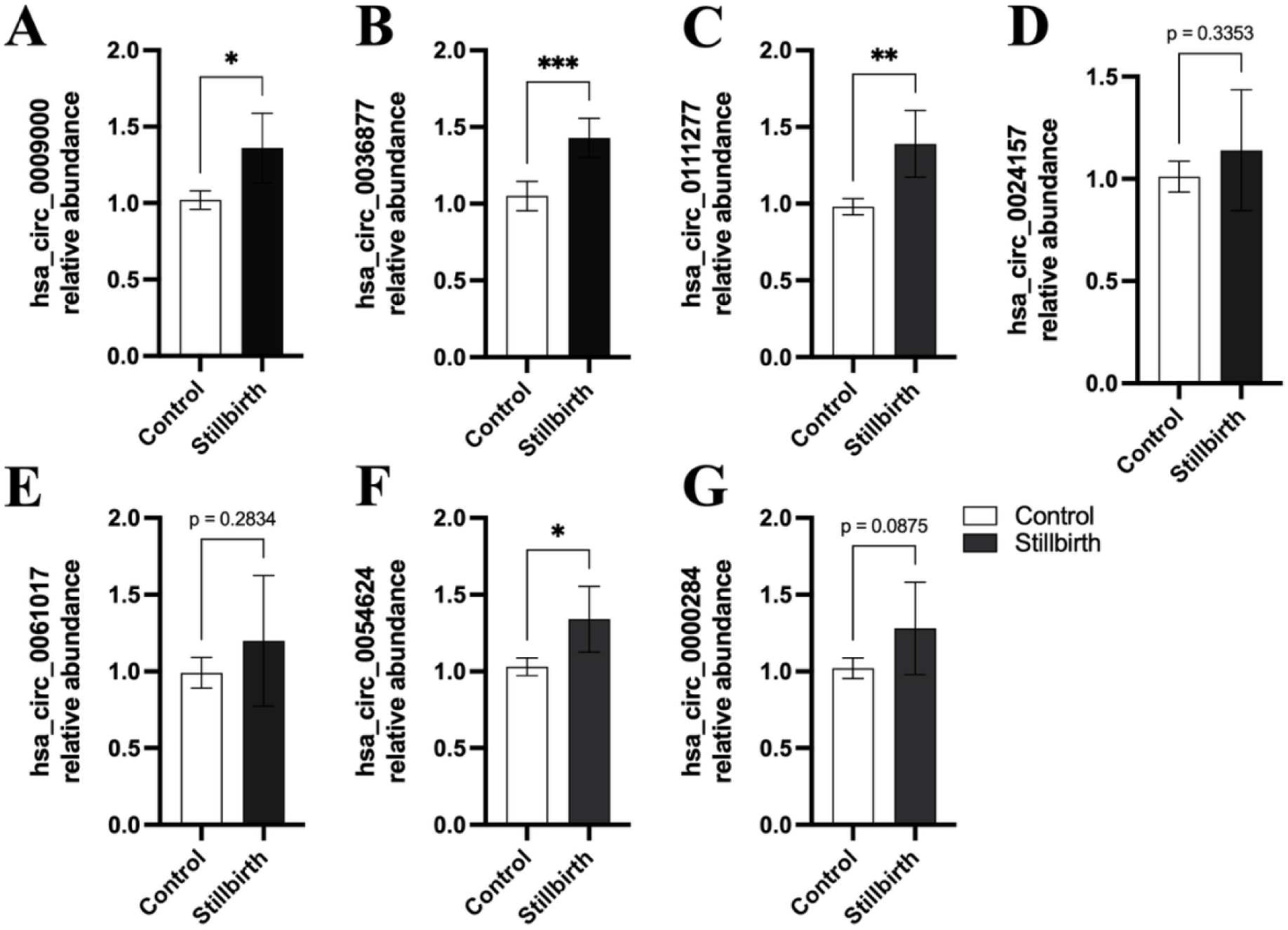
Abundance (as fold change relative to housekeeping genes) of **(A)** hsa_circ_0009000, **(B)** hsa_circ_0036877, **(C)** hsa_circ_0111277, **(D)** hsa_circ_0024157, **(E)** hsa_circ_0061017, **(F)** hsa_circ_0054624, **(G)** hsa_circ_0000284 in maternal blood sampled at 15-16 weeks’ gestation from women who went on to have a stillbirth and women who had live births. n = 12 / live births, n = 6 / stillbirth group. Data are presented as mean ± standard deviation. * p < 0.05, ** p < 0.01, *** p < 0.001. Results are considered statistically significant if p < 0.05.

## 4. Comment

### Principal Findings

Here, we confirm that placentae from stillbirth experience accelerated ageing (as determined by typical hallmarks of decreased telomere length, increased rates of cellular senescence and DNA damage). circRNAs circ_0009000, circ_0024157, circ_0061017, circ_0036877, circ_0054624, circ_0111277 and circ_0000284 accumulate in ageing placental tissue, and prematurely accumulate in stillbirth placentae. These circRNAs bind placenta DNA, facilitating DNA breaks and cellular senescence. Measurement of candidate circRNAs in maternal blood sampled at 15-16 weeks’ gestation is a potential screening tool to indicate premature placental ageing indicative of stillbirth risk.

## Results in the Context of What is Known

### The accumulation of circRNAs

The accumulation of circRNAs in aged tissue is a relatively recent discovery, evident from studies^35^ using *Drosophila melanogaster* photoreceptor neurons^36^, mouse brain and heart^8^ and *Caenorhabditis elegans*^7^. It is likely that circRNAs accumulate in tissues as their lack of a 5’ cap and polyadenylated tail renders them resistant to exoribonuclease degradation^37^. Another explanation for circRNA accumulation in some tissues is that in cell types such as neurons, rates of mitosis and proliferation are lower than other cell types, allowing reduced turnover of circRNAs^35,38–40^. Given that the nuclei of the syncytiotrophoblast are non-mitotic^41^, transcription is significantly reduced^42^ and the syncytiotrophoblast displays markers of senescence^43,44^, this could be a primary area of circRNA accumulation. Indeed, this is consistent with our *in vitro* findings of circRNAs expression primarily in the syncytiotrophoblast.

Recently, circRNAs were identified as a potential forensic tool for estimating biological age in human blood^45^, which is an exciting leap from circRNA accumulation in animals to differential circRNA expression with age in humans. Other studies have shown differentially expressed circRNAs in human granulosa cells from women who were young versus those of advanced reproductive age^46^, and that specific circRNAs in human peripheral blood were associated with parental longevity and cellular senescence^47^. The link between circRNA differential expression and Alzheimer’s disease is becoming more widely investigated^48^, which is of particular interest to this study given that Alzheimer’s disease is, at its core, an ageing disorder^49^. Whilst these studies demonstrate that expression of circRNAs is *changed* between differently aged groups, none of these studies investigated circRNAs *accumulation* in ageing tissue.

We posit that the abundance of a circRNA is a combination of expression level as well as accumulation. circRNA expression level can be increased by overexpressing the linear gene, or by alternative splicing that promotes the circular over the linear transcript(s). circRNA accumulation indicates that the circRNA remains in the tissue^35^, unaffected by exoribonucleases^37^. Over time, the abundance of the circRNA in the tissue increases as the accumulated transcripts escape degradation. This occurs without change in the linear gene expression nor alternative splicing. The result is that these circRNAs can continually exert their function in the tissue well after their linear gene’s expression and at higher-than-expected abundance.

### Implications of circRNA accumulation as a novel mechanism for placental ageing

Like other tissues in the body, the placenta undergoes the normal phenomenon of ageing. Tissue ageing includes progressive increases in cellular senescence^50^. For the placenta, this implies a decline in functionality as gestational age increases, resulting in a reduced capacity of the placenta to support the fetus^3^. Placentae from stillbirths undergo an accelerated ageing process^3^, exhibiting similar pathological features regardless of their earlier gestational ages. Markers of ageing such as increased DNA and RNA oxidation, lipid peroxidation, altered perinuclear location of lysosomes and larger autophagosomes are found in placentae from stillbirths as well as those from healthy late term compared with placentae from healthy early term^3^. This study expands on the current literature, demonstrating shorter telomeres, increased cellular senescence and DNA breaks occur with placental ageing.

A circR-loop is formed when a circRNA molecule binds one strand of helicase-unwound genomic DNA^10^. The steric hindrance of this bulky structure causes RNA polymerase II to stall during transcription, resulting in a DNA break, and often a subsequent fusion at another locus^51^, promoting genomic instability^11^. Double-stranded DNA breaks are lethal to cells, as they affect both strands of DNA and promote the loss of genetic information^13^. Any type of DNA damage can trigger cellular apoptosis or senescence^12^. Given that we observed increased DNA breaks in stillbirth placentae, which also had elevated abundance of circ_0000284 binding to placental DNA, we further determined that circ_0000284 induces DNA damage and cellular senescence *in vitro*. Moreover, RNase H1 was investigated as a proposed method of circRNA degradation. Knockdown of RNase H1 using siRNA revealed that a portion of all circRNAs are susceptible to RNase H1 degradation in placenta tissue, however this was more impactful in placentae from unexplained stillbirth. This could be due to a greater proportion of circRNAs susceptible to RNase H1 in tissue, compared to circRNAs that are not susceptible, or could indicate a lack of endogenous H1 in placentae from unexplained stillbirth. This would also potentially account for the premature circRNA accumulation in these samples. Together, this presents a promising avenue for investigation as a mechanism for stillbirth via premature placental ageing. This study also provides evidence that circRNAs accumulate in healthy ageing placental tissue, and can also be detected as a novel proxy measure of placental ageing in maternal blood.

### Clinical Implications

Our group 41+ weeks’ gestation encompassed women who birthed between 41+1 and 41+6 weeks’. Given that gestation of >40 weeks, in addition to maternal age ≥35 years^3^ and male fetal sex^1^, are risk factors for stillbirth, the current Pregnancy Care Guidelines consider 42 weeks’ and beyond as post-term and recommend fetal monitoring and induction of labour from 41 weeks’, even in uncomplicated, low risk patients^52^. Indeed, evidence shows that stillbirth and neonatal death can be reduced with fetal surveillance twice weekly from 39 weeks’^53^.

In this study, placentae from stillbirths displayed evidence of further ageing hallmarks. As these stillbirth placentae were collected at gestational weeks 23, 26, 31 and 34, this clearly indicates that in stillbirth the placenta experiences accelerated tissue ageing, resulting in a post-term phenotype much earlier than would be appropriate for their gestational age. Use of a blood test for signalling accelerated placental ageing, as supported by our results, could be beneficial to screen for pregnancies at high risk of stillbirth.

### Research Implications

Interpreting our results, we propose the following mechanism as a potential contributor to stillbirth pathology (**Figure 5**): In uncomplicated pregnancy, circRNAs naturally accumulate in placental cells as the tissue ages with gestation. However, in pregnancies resulting in unexplained stillbirth, premature placental ageing results in accelerated accumulation of circRNAs. These circRNAs form circR-loops, with at least one circRNA (circ_0000284) known to facilitate DNA damage and induce senescence. As DNA breaks facilitate genomic instability^11^ and cellular senescence^54^ and apoptosis^55^, this may result in reducing placental function to the detriment of the fetus.

**Figure 5.**
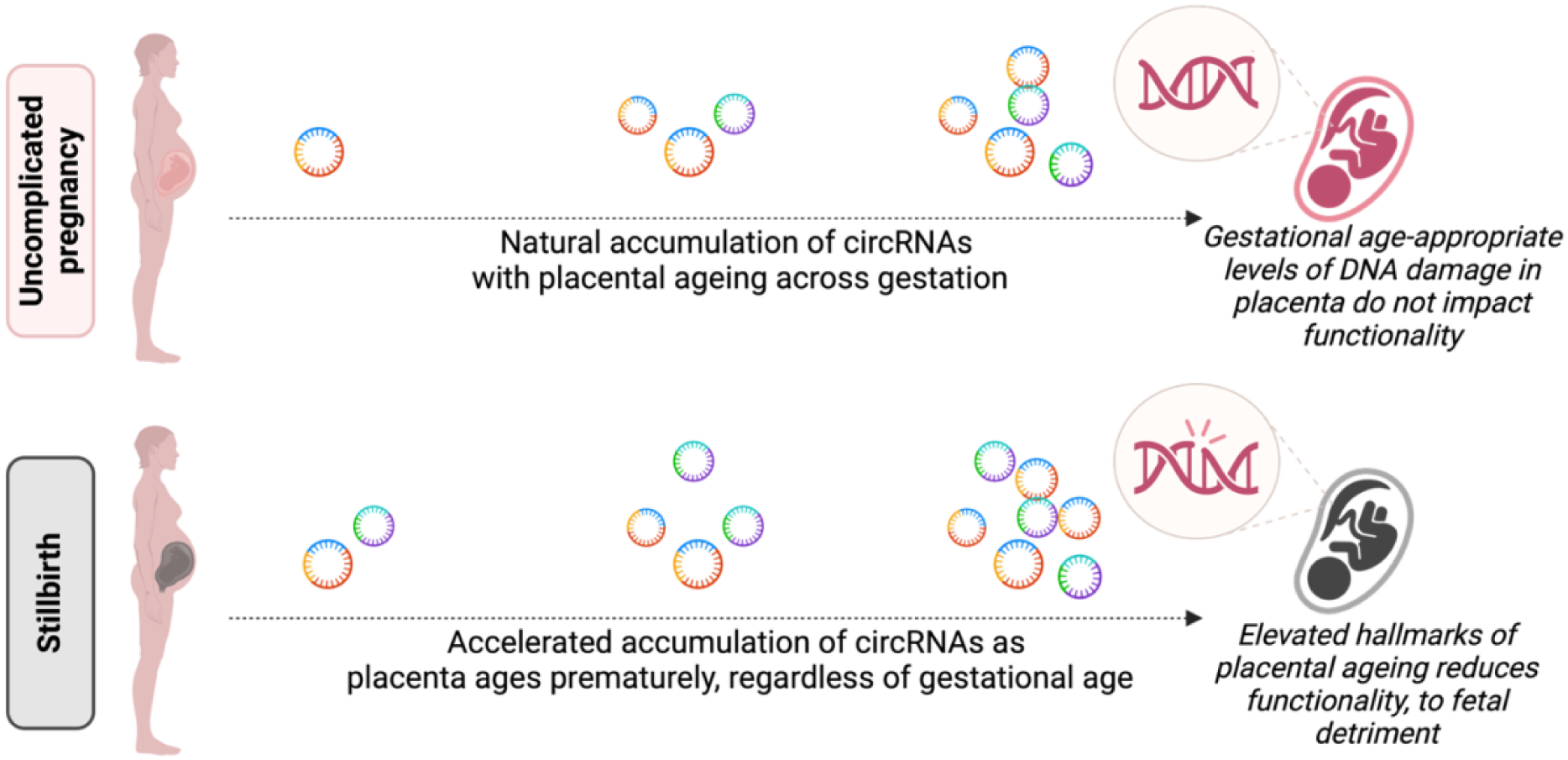
Proposed mechanism by which circRNA accumulation and placental ageing contribute to stillbirth.

### Strengths and Limitations

We confirm that circRNAs accumulate in human aged tissue. Importantly, we established that there is no global increase in total RNA, nor circular RNA, abundance with placental gestational age; hence, our candidate circRNAs are unique in their ability to accumulate with age. It should be noted that accumulation of these specific circRNAs with age may be placenta-specific, as this study only examined placental tissue, and this candidate list may not be exhaustive. Further studies will profile all accumulating circRNAs in the placenta as it ages.

Additionally, this study is limited by the small sample size of stillbirth placentae; however, this is a common issue in stillbirth research^3,56^ given the difficulty in obtaining tissue within an appropriate timeframe. As stillbirths that could be explained by fetal congenital abnormality were excluded, resulting in a pure “unexplained stillbirth” sample pool, this further reduced available tissue samples. Moreover, sample size limited our ability to perform fetal sex-specific statistical tests. Further studies will be conducted with larger sample sizes to allow for increased comparisons. Future studies will also include participants of different ethnicities, accounting for ethnicities which are more prone to stillbirths (such as South Asian women^53^).

## Conclusions

circRNAs accumulate in ageing human placental tissue, in the absence of any change in total linear or circular RNA expression nor alternative splicing variants. Importantly, circRNAs circ_0009000, circ_0024157, circ_0061017, circ_0036877, circ_0054624, circ_0111277 and circ_0000284 are present at exceptionally high levels in placenta from stillbirths, similar to or surpassing the circRNA levels found in post-term placentae. We provide evidence that these circRNAs bind to DNA to form circR-loops in placental tissue. *In vitro* experiments in primary human syncytiotrophoblasts show that knockdown of circ_000284 results in decreased DNA breaks in cells, indicating that this circRNA facilitates DNA damage, and decreased expression of senescence-associated genes, implying that this circRNA induces cellular senescence. Finally, measurement of candidate circRNAs in maternal blood sampled at 15-16 weeks’ gestation is a viable tool to screen for accelerated placental ageing, as a proxy for risk of stillbirth.

## Acknowledgments

We thank the women who consented to their placental tissue being used for this research. Thank you to the clinical and anatomical staff from the Women’s and Children’s Hospital and Lyell McEwin Hospital who were involved in the women’s care, and who collected and biopsied the placental samples. Thank you to Professor Jose Polo for his insightful suggestions. Figures were created using Biorender.com

## Author Contributions

A.L.A. conceptualised this work, performed experimental work and formal analysis of data, original draft preparation, and review and editing. M.R.J., H.S.S. and C.P.B. provided stillbirth placenta samples and reviewed and edited the manuscript. D.M. contributed to experimental work and data analysis. S.W. provided molecular expertise and reviewed and edited the manuscript. M.D.S. and T.J-K. reviewed and edited the manuscript. G.A.D. provided clinical expertise and reviewed and edited the manuscript. C.T.R. provided supervision, funding acquisition, interpretation of data, review and editing of the manuscript. All authors contributed to the article and approved the final version.

---

**Supplementary Figure 1**. (A-B)RNA concentration (ng / µL) and **(C)** abundance (as fold change of RNase R-depleted RNA relative to total RNA) in isolated cells from 37, 38, 39, 40 and 41+ uncomplicated, and stillbirth (S), placentae. n = 12 placentae / uncomplicated gestational group, n = 4 stillbirth placentae. RNA was extracted from each sample, then split into two aliquots; the first was untreated (total RNA) while the second was digested with RNase R (leaving only circRNAs). RNA concentration and abundance of total RNA and RNase R-digested RNA were assessed to determine whether these changed across gestation or in stillbirth. Data are presented as scatter plots with bars and indicate mean ± standard error of the mean. ns = not significant.

**Supplementary Figure 2**. Abundance (as fold change relative to abundance at 37 weeks’ gestation) of linear CRHR2 gene variants **(A)** XM_024446665.1, **(B)** XM_017011752.2, **(C)** NM_001202483.2, **(D)** NM_001202482.2, **(E)** NM_001202481.1 and **(F)** NM_001202475.1; linear CWF19L2 gene variants **(G)** XM_017017239.2, **(H)** XR_947795.2, **(I)** NM_152434.3 and **(J)** XM_011542620.3; linear RAB22A gene variant **(K)** NM_033109.5; linear FURIN gene variants **(L)** NM_001382622.1, **(M)** NM_002569.4, **(N)** NM_001382619.1, **(O)** NM_001289823.2, **(P)** NR_168464.1, **(Q)** NM_001382620.1, **(R)** NM_001382621.1 and **(S)** NM_001289824.2; linear PNPT1 gene variant **(T)** NM_033109.5; linear PAPPA2 gene variant **(U)** XM_005245422.3; and linear HIPK3 gene variants **(V)** NM_001278163.2, **(W)** NM_001048200.3, **(X)** NM_005734.5, **(Y)** XM_04746224.1, **(Z)** NM_001278162.2, **(AA)** XM_047426223.1 and **(AB)** XM_047426225.1 all in isolated cells from 37, 38, 39, 40 and 41+ uncomplicated, and stillbirth, placentae. n = 12 placentae / uncomplicated gestational group, n = 4 stillbirth placentae. Data are presented as scatter plots with bars, and indicate mean ± standard error of the mean. ns = not significant.

