## Supplementary figures and images for "Circular RNAs accumulate in ageing human placental tissue and in stillbirth, leading to DNA damage and cellular senescence"

### Supplementary Figure 1

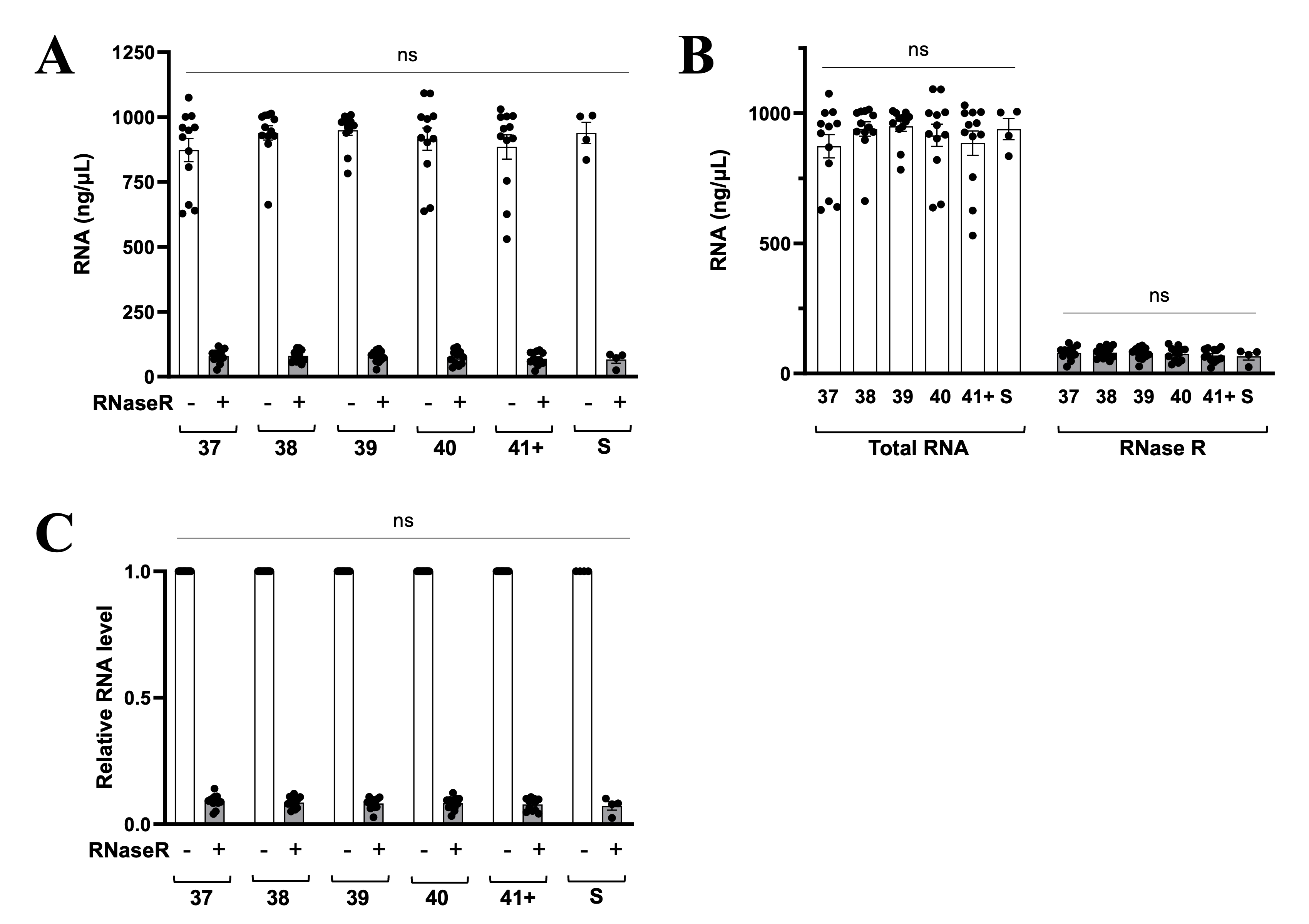
